# Modular response analysis reformulated as a multilinear regression problem

**DOI:** 10.1101/2022.08.11.503572

**Authors:** Jean-Pierre Borg, Jacques Colinge, Patrice Ravel

## Abstract

**Motivation:** Modular response analysis (MRA) is a well-established method to infer biological networks from perturbation data. Classically, MRA requires the solution of a linear system and results are sensitive to noise in the data and perturbation intensities. Applications to networks of 10 nodes or more are difficult due to noise propagation.

**Results:** We propose a new formulation of MRA as a multilinear regression problem. This enables to integrate all the replicates and potential, additional perturbations in a larger, over determined and more stable system of equations. More relevant confidence intervals on network parameters can be obtained and we show competitive performance for networks of size up to 100. Prior knowledge integration in the form of known null edges further improves these results.

**Availability and implementation:** The R code used to obtain the presented results is available from GitHub: https://github.com/J-P-Borg/BioInformatics

## Introduction

Biological systems are orchestrated by a multitude of interaction networks. At a cellular scale, the knowledge of protein physical interaction networks, phosphorylation cascades, or gene regulatory networks is central in our understanding of homeostasis, disease, reaction to environmental changes, and many other situations. The inference of biological networks from experimental data has thus attracted much attention from the computational biology community (Huynh-Thu and Sanguinetti, 2019).

A common experimental design to learn about the connectivity between a set of molecules of interest is to measure their activity under different conditions. Molecules can be for instance genes, transcripts, proteins, or metabolites, and the notion of activity might relate to their abundance or state, *e*.*g*., the phosphorylation level of a protein. Modular response analysis (MRA) is a method tailored for a particular experimental design: data obtained through the systematic perturbation of each molecule activity under the assumption that the biological system has reached a steady state at the moment of the measure (Kholodenko *et al*., 2002). Under these hypotheses, MRA is a versatile framework allowing to infer connectivity between molecules (or groups of molecules called modules) *X*_1_, …,*X*_*N*_ simply assuming that they are related by an equation of the form 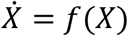. The knowledge of *f* is not necessary because the steady state hypothesis combined with the implicit function theorem enable solving a linearized problem based on experimental data only. Except for the steady state hypothesis, which is not fulfilled by every system obviously, MRA provides an elegant solution involving linear algebra only to explore biological networks quantitatively. It has been applied and extended by a number of researchers (Santra *et al*., 2018; Dorel *et al*., 2018; Jimenez-Dominguez *et al*., 2021; Mekedem *et al*., 2021). However, MRA is sensitive to measurement noise and to the intensity of the perturbations exerted on network nodes (Andrec *et al*., 2005; Thomaseth *et al*., 2018). Different approaches have been suggested to alleviate these problems (Santra *et al*., 2018), *e*.*g*., to perform a bootstrap followed by an estimation of confidence intervals (CIs) of the connectivity coefficients (Santos *et al*., 2007; Thomaseth *et al*., 2018; Jimenez-Dominguez *et al*., 2021).

In this report, we combine MRA with multilinear regression. Indeed, MRA equations can be interpreted as a linear regression problem (see below) thus enabling the use of any linear regression strategy. This new formulation hence defines a family of methods, which we will illustrate employing classical algorithms such as the least square, LASSO, or STEP. One advantage of the regression approach is to integrate the treatment of the experimental noise with the model accuracy estimation in terms of residual variance. That is, how well the linear MRA model approximate the potentially nonlinear biological system and how accurate are the estimated network parameters. Using data from DREAM 4 Challenge, we show that we can apply MRA modeling to 10-to 100-node networks with competitive performance, while a standard resolution of MRA equations already encounters difficulties with 10-node networks.

## Methods

### Fundamental definitions

We consider a putative biological network comprised of *N* modules that potentially interact. A module can be a molecule (transcript, protein, …) or it can represent the contribution to the network of a set of molecules such as a pathway or part of a pathway. In the latter case, one measurement reports for the activity of the whole module. MRA aims at determining signed connectivity coefficients between the modules. The module activity levels are represented as a time-dependent function *X*: ℝ^+^ ↦ ℝ^*N*^. In case the modules were gene transcripts, *X*(*t*) could for instance represent their abundance over time. We assume the existence of a continuously differentiable function *f*: ℝ^*N*^× ℝ^*M*^ ⟼ ℝ^*N*^and a vector of intrinsic parameters *P* ∈ ℝ^*M*^, at least one *per* module (hence *M* ≥ *N*), such that

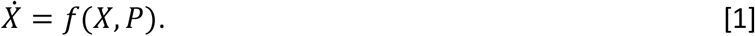

The function *f* is usually unknown. *P*_0_ represents the biological system in its basal state, *i*.*e*., in the absence of any perturbation. We suppose the existence of a time *t*_0_ after which the system has reached a steady state (including all the perturbed states):

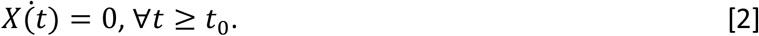

Let *X* = *X*(*P*_0_) be a solution of the system [2]. The application of the implicit function theorem gives us an exact formula for the connectivity coefficient representing the influence of a module *j* on another module *i* (Kholodenko *et al*., 2002; Jimenez-Dominguez *et al*., 2021):

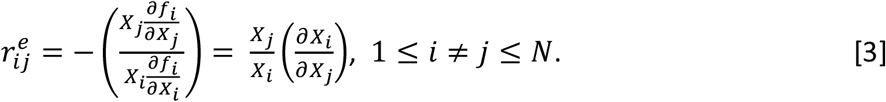

Note that 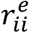 is not defined by Eq. [3]. By convention, we set 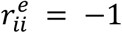, see (Jimenez-Dominguez *et al*., 2021) for an algebraic justification.

Taylor’s development at the first order enables us to write an expression for the system under perturbation Δ*P* (Jimenez-Dominguez *et al*., 2021).

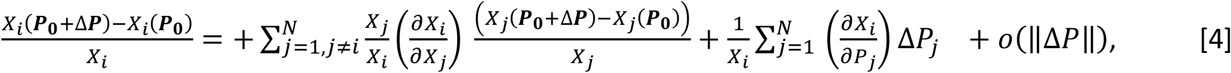

### Standard MRA formulation

Only one perturbation *per* module is considered implying *M* = *N*. Experimental measurements obtained under perturbation *q*_*i*_ result from the modification of the intrinsic parameter *P*_*i*_ that directly affects node *i* only. Accordingly,

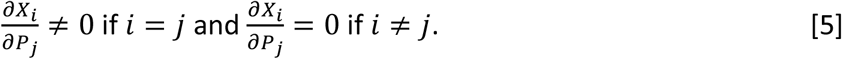

We introduce the notation

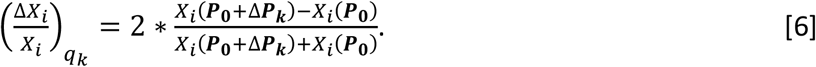

Using the mid-point in the denominator of Eq. [6] right-term instead of *X*_*i*_(***P***_**0**_) is customary to avoid divisions by 0. Thanks to Eq. [4] and [5], we obtain the system of equations

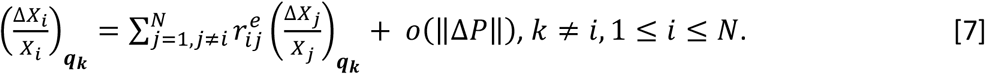

Ignoring the error term *o*(∥Δ*P*∥) in [4] that represents departure from linearity, we obtain its linear approximation

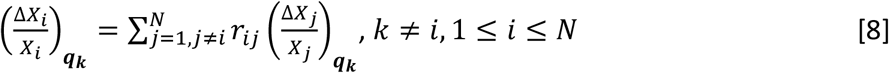

that is solved by MRA (note that *r*_*ij*_ replaces the exact 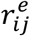). In particular, writing matrices *r* = (*r*_*ij*_) and 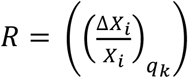, the connectivity coefficient are obtained by:

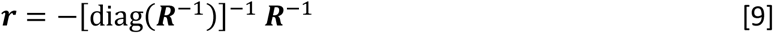

an *N*× *N* system (Kholodenko *et al*., 2002).

### A multilinear regression formulation

Eq. [9] does not take into account the existence of replicates in the data. It requires a single value for each perturbed state and each module. In practice, *ad hoc* procedures must be added to exploit replicates properly and estimate confidence intervals for the *r*_*ij*_ computed after Eq. [9]. Typically, bootstrap approaches are used that apply Eq. [9] to resampled data or Eq. [9] is applied to averaged data perturbed by an appropriate noise function (Andrec *et al*., 2005; Santos *et al*., 2007; Thomaseth *et al*., 2018; Jimenez-Dominguez *et al*., 2021).

An alternative approach is to consider Eq. [8] as the homogeneous multilinear regression of 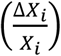 with regression parameters *r*_*ij*_. In that case, the system to solve is not limited to *N* − 1 equations. We can have one equation *per* replicate with any number of replicates. It is even possible to integrate multiple perturbations at each node, *e*.*g*., at different intensities or by different means, and hence *M* ≥ *N*. Obviously, we have *N*such regressions to perform (*N* − 1 variables each) and the regression approach let us estimate *r*_*ij*_ variability, directly taking advantage of all the tools provided by the regression literature (Hastie *et al*., 2009).

### Considered regression methods

To solve Eq. [8] with a regression approach leaves the choice of the specific regression method free. Here, we considered several standard methods. An obvious choice was least square estimation (LSE) applied to multilinear regression. Because most biological networks are rather sparse and they can reach a certain size, we primarily considered methods able to eliminate the least significant regression parameters. Accordingly, we defined the LSE_CI method as the application of LSE followed by an elimination of all the *r*_*ij*_ whose 95% CI included 0.

A second method was threshold linear regression (TLR), which also starts with LSE but eliminates coefficients differently. The rule is to compute a threshold *T*_h_ as the 25^th^ percentile of all the estimated values |*r*_*ij*_| and to impose:

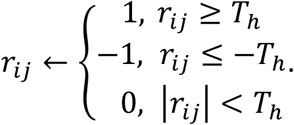

The third method was LASSO, a shrinkage estimator. Using LASSO for a node *i* (the i^th^ of the *N*systems of size *N* − 1 to solve) is defined by :

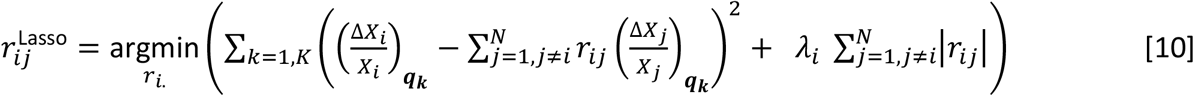

The choice of the hyper-parameter *λ*_*i*_ is determined by cross-validation (CV) (Hastie *et al*., 2009) and it is hence necessary to have experimental replicates or multiple perturbations at certain nodes.

The fourth option was methods integrating a variable selection scheme. Namely, we used STEP forward (STEP-Fo), STEP backward (STEP-Ba), and Stepwise regression (STEP-Bo) combining both backward and forward (Hastie *et al*., 2009). They essentially consist in finding subsets of variables minimizing the sum of the residual squares while ignoring the other variables according to different selection strategies. This selection process introduces a bias in the solution, but here this bias should remain modest because the networks are sparse (roughly 80% of null edges).

### Prior knowledge integration

Let us assume that we know the null edges of node *X*_*i*_, *i*.*e*., the set *A*_*i*_ = {*j* ∈ {1, …,*N*}: *r*_*ij*_ = 0}. Each index *j* ∈ *A*_*i*_ cancels a column of the linear system associated with solving node *X*_*i*_ connectivity coefficients. When the regressive approach is used, Eq. [8] becomes:

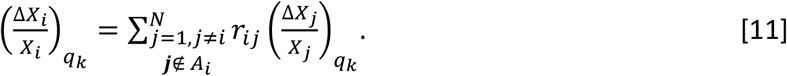

Hence, the linear system corresponding to node *X*_*i*_ is reduced to *N* − 1 − card(*A*_*i*_) parameters to estimate. The number of degrees of freedom of the model and its residual variance decrease accordingly.

## Results and Discussion

### Impact of noise and perturbation intensities on MRA solutions

Conceptually, the derivation of MRA equations entails infinitesimal perturbations of the network modules (Kholodenko *et al*., 2002) such that classical tools of calculus can be applied, notably the implicit function theorem that is a local result (Jimenez-Dominguez *et al*., 2021). In practice, infinitesimal perturbations are usually not feasible and not advisable because experimental data tend to be noisy. Nevertheless, Taylor’s development in Eq. [4] shows a dependence of the error on the perturbation intensity (error term) underlying the potential loss of accuracy of MRA depending on the strengths of the variable nonlinear dependencies. Accordingly, we decided to study the relationship between accuracy, noise, and perturbation strengths on a small but representative system. We used the synthetic kinases network (MAP, MKK, MKKK) introduced by Kholodenko (Kholodenko *et al*., 2002) in MRA original paper. This system is comprised of six molecules (counting the phosphorylation states) and, unlike Kholodenko’s analysis that was limited to three modules, we considered the whole system with six modules, one *per* molecule (Fig. 1a). The dynamical equations are given in Supplementary Information (SI). We solved them numerically using the R package ode and the time *t*_0_ after which a steady state is reached was determined by a procedure detailed in SI as well. The exact connectivity coefficients (Fig. 1b) were computed numerically.

**Figure 1.**
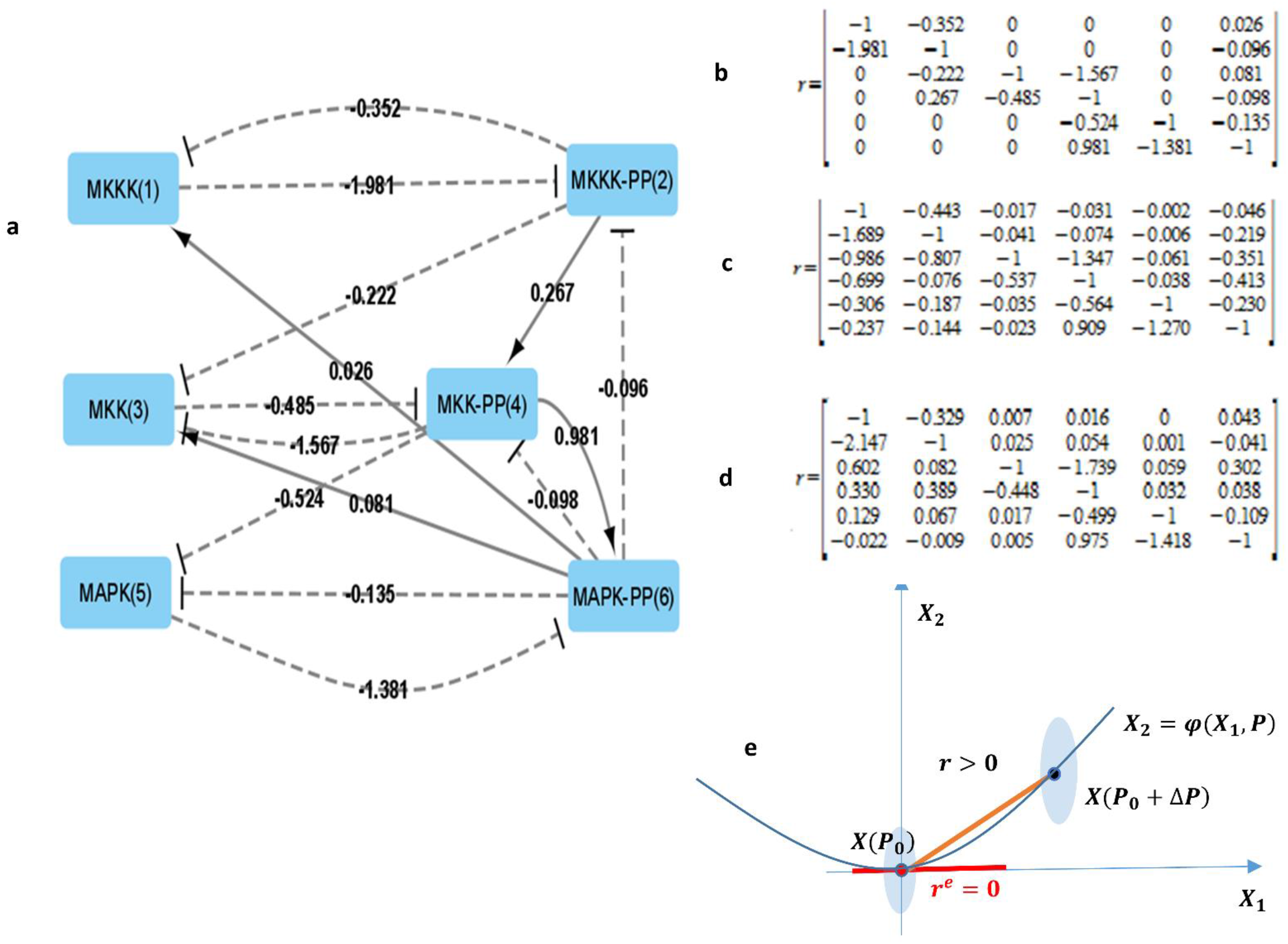
A 6-node MAP kinase network. (**a**) Exact network topology with connectivity coefficients 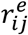. Node names correspond to the different forms of kinases, and the numbers in brackets are their identifier in further figures. A solid arrow means a direct activation, while a dashed arrow means an inhibition. (**b**) The matrix of exact 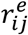. (**c**) Estimated *r*_*ij*_ under -50% perturbation. (**d**) Estimated *r*_*ij*_ under +50% perturbation.(**e**) Illustration of the impact of nonlinearity on the null edges problem. The exact connectivity coefficient *r*^*e*^ = 0 (slope of the implicit function *φ* on *X*(*P*_0_)) is approximated by the slope *r* > 0 of the segment (*X*(*P*_0_), *X*(*P*_0_ + Δ*P*)) because of Taylor’s development restricted to the first order.

We started with data devoid of noise and perturbations *q*_*k*_ applied to the six speed constants of the system 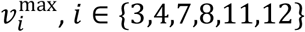, *i* ∈ {3,4,7,8,11,12} (see SI) and Δ*P*_*k*_ equals to ± 50 %. Such perturbations could be induced by kinase inhibitors for instance. For each perturbation, a new steady state *X*(*P*_0_ + Δ*P*_*k*_) was computed. A first observation was that due to nonlinearity, symmetrical perturbations did not result in symmetrical differences compared to the basal state (Fig. 1cd). In particular, the error was greater with −50% perturbations than with 50% perturbations, and was a nonlinear function of ∥Δ*P*∥ (Fig. 2a). Moreover, we observed that connectivity coefficients that were zero in the basal state, were no more zero when estimated with standard MRA (Fig. 1b-d), though MRA is meant to estimate their basal state value. Departure from zero could indeed be substantial in such a small network, which is a limitation for identifying the topology of biological sparse networks (Fig. 1e).

**Figure 2.**
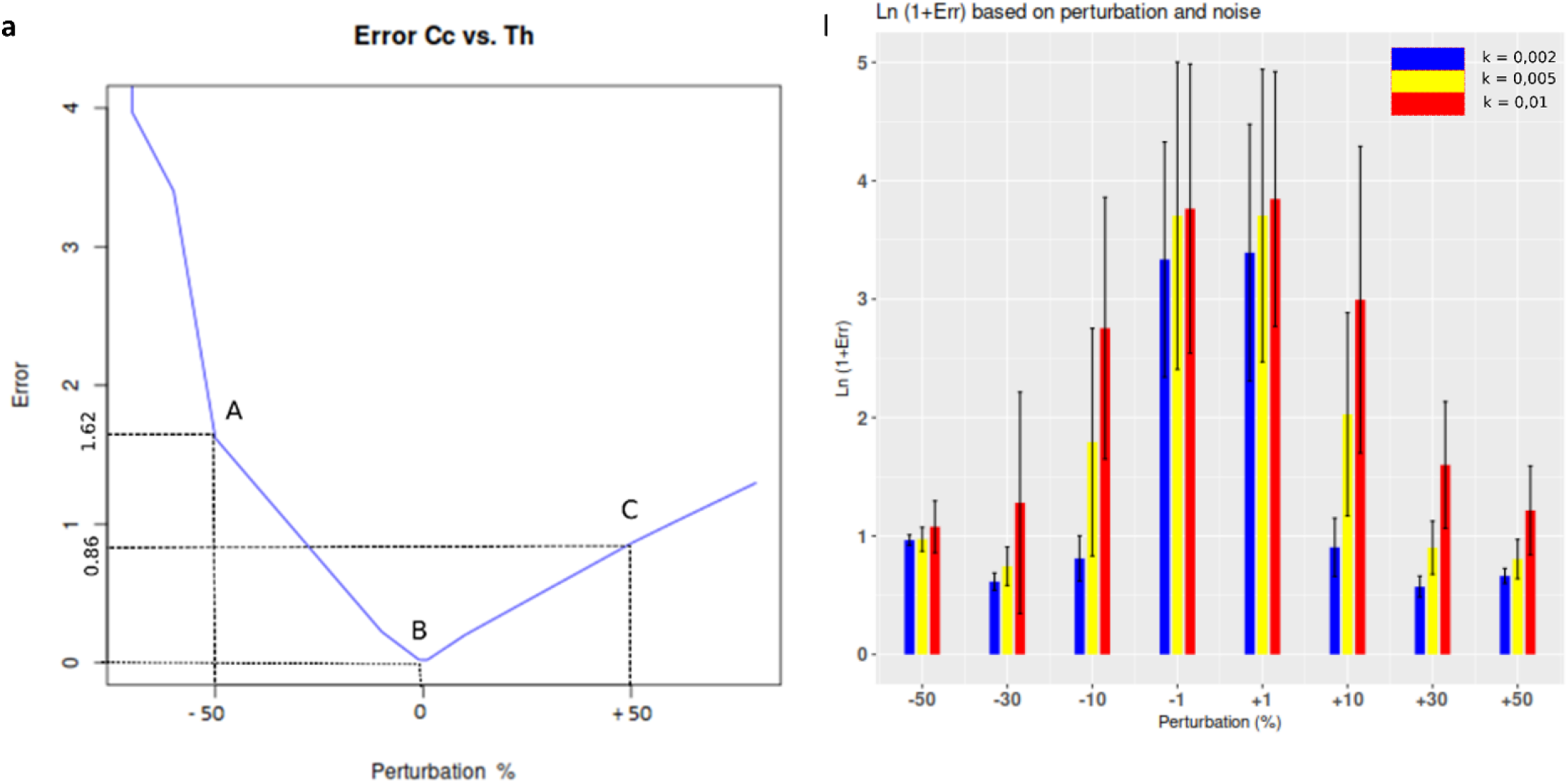
Impacts of perturbation and noise level for the 6 MAP kinase network. (**a**) The squared error 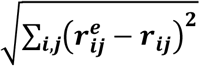 as a function of the perturbation intensity in the absence of noise. We note that points A and C are not symmetrical with respect to B although symmetrical perturbations were applied. (**b**) The impact of perturbations at different intensities in the presence of Gaussian noise at 0.1%, 0.5%, and 1%. Black vertical lines indicate the error standard deviation. We note that the error is huge with small perturbations (-1% and +1%) applied to noisier data. Stronger perturbations yield smaller errors provided the noise remains reasonable. Otherwise, errors grow rapidly as well (vertical axis in log).

Naively, one would address the issue above by using smaller perturbations, but as already mentioned, in the presence of noise, this approach might be inapplicable. Three levels of Gaussian additive noises *N*(0, *σ*) were added to the *X*_*i*_ with 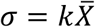, where 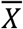 was the mean of all the theoretical concentrations at steady state and *k* was set at 0.1%, 0.5%, or 1%. Note that with *k*=1%, a substantial noise is added to low signals due to 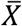 definition. The error in the presence of noise (Fig. 2b) behaved in a reverse fashion compared to what we observed without noise. This is because the noise error was larger than the nonlinearity error 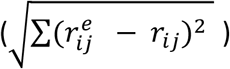. With more noise it is indeed necessary to increase perturbation intensities to obtain a sufficient signal-to-noise ratio, but this also drives us away from the exact solution in case the system is substantially nonlinear with respect to *P* (Eq. [4]). In what follows, we used +50% perturbations as a reasonable compromise.

### Determining edges existence

MRA assesses network edges quantitatively, but measurement noise usually generates nonzero values for all the connectivity coefficients. Figs 3a and 3b illustrate this phenomenon for 6-node kinase network introduced above. For standard MRA, we estimated 95% CIs applying a bootstrap with 100 repetitions for two noise levels (0.1% and 0.5%). A connectivity matrix *r* is obtained for each repetition of the bootstrap and CIs were simply obtained as the 2.5^th^ and 97.5^th^ percentiles. We observed an already known relationship (Thomaseth *et al*., 2018): CI sizes increase with higher noise in the data, independent of the number of replicates. It is important to note that such CIs represent the consequence of the total variance and by applying the standard MRA approach we are limited to such variance estimations *a posteriori, e*.*g*., via simulations such as the bootstrap. The independence on the number of replicates is an obvious limitation. On the contrary, solving MRA with multilinear regression methods enables to estimate the residual variance, *i*.*e*., the variance caused by the nonlinear contributions. Residual variance-based CIs is common practice when applying linear models to data. In Figs 3a and 3b, we show such CIs for 3 and 5 replicates and noise levels at 0.1% and 0.5% for LSE regression. Residual variance-based CIs were indeed strongly reduced compared to standard MRA total variance-based CIs, and more replicates yielded smaller CIs. In a previous study limited to 3-module networks (Thomaseth *et al*., 2018), the idea of applying LSE has been already envisioned. Nonetheless, LSE was applied to each replicate separately thus leading to poor performance. In one case, the authors linked dual perturbations and reported encouraging performance, but they did not follow up on this observation.

**Figure 3.**
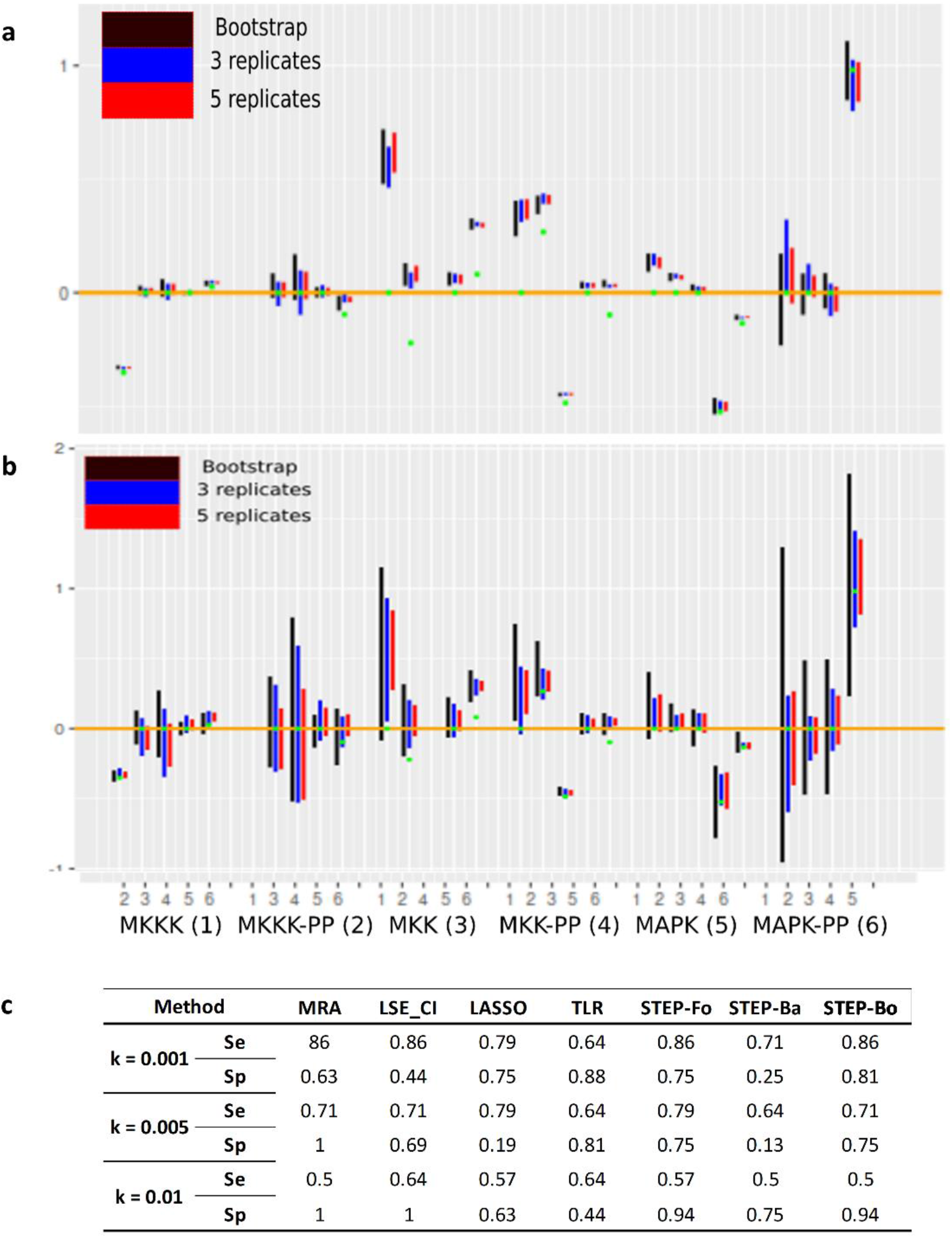
Inference performance for the 6 MAP kinase network. (**a**) Estimated connectivity coefficient *r*_*ij*_ 95% CI for MRA standard resolution followed by a bootstrap, LSE with 3 or 5 replicates and Gaussian additive noise at 0.1%. Green dots represent the exact values. (**b**) Same as (a) but with 0.5% noise. The x-axis indicates the edge origins, *e*.*g*., MKKK, and the numbers the edge targets (**c**) Performance estimations based on edge directionality and existence or absence. Se = sensitivity and Sp = specificity.

Regarding edge existence, we reduced the exact connectivity coefficient 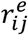 to -1, 0, or 1 to simplify performance comparisons. With the estimated *r*_*ij*_, we applied the criterion described above for the LSE_CI method: an edge is deemed significant and hence non-null if 0 is excluded from its connectivity coefficient CI, and assigned +1 or -1 depending on whether 0 is on the left or on the right of the CI. Results comparing standard MRA to various MRA-regression combinations are reported in Fig. 3c and Fig. S1. We see that on this modest 6-node network, MRA and MRA-regression performance depends on the noise level. Overall, the best sensitivity was achieved by STEP-Fo, LSE_CI, and LASSO (in this order), which outperformed MRA. TLR remained stable, regardless of the noise. Concerning specificity, the best results were obtained by MRA, STEP-Fo, and STEP-Bo. Upon noise increase, specificity was improved for MRA and LSE_CI because an increased noise inflated the CIs leading to the correct detection of every null edge (Suppl. Table 1).

### Medium size networks from DREAM 4 Challenge

The DREAM 4 Challenge released five 10-gene and five 100-gene networks of *Escherichia coli* meant to compare inference algorithm performance. Each gene was subjected to two independent perturbations series, knockdowns (KDs) and knockouts (KOs). All the gene expression levels *X*_*i*_ were normalized and an unknown noise was added. The exact solutions were provided as binary edges, *i*.*e*., *r*_*ij*_ ∈ {0,1}. The networks were sparse with 80% of null edges on average. DREAM 4 data regarding the network dynamics before reaching a steady state were ignored by us. We evaluated the performance of our methods on the existence or nonexistence of the inferred edges. In the case of MRA, it was not possible to combine KO and KD data in one analysis. We hence treated the two sets of perturbations separately.

Figure 4a features the first 10-gene network compared with the result found by TLR. Performances of the various methods are reported in Fig. 4b averaging over the five networks of each size. On these larger (10 genes) and much larger (100 genes) networks, regressive approaches showed strong superiority over standard MRA. MRA-KD is close to random selection for 10- and 100-gene networks, whereas MRA-KO performs reasonably with 10-gene networks but is also close to random with 100 genes (Suppl. Figs 1-3). The simple regression methods LSE_CI and TLR delivered better performance, but were nonetheless sensitive to the network size. Regression methods adapted to sparse solutions (STEP-xx and LASSO) performed robustly and STEP-xx provided the best compromise overall. Comparing these methods with DREAM 4 Challenge ranking established in 2014, based on P-values concerning areas under the ROC curve (official criterion), we found that STEP-xx and TLR methods ranked 3^rd^ for 10-gene networks. They respectively ranked 1^st^ and 8^th^ for 100-gene networks (STEP-Fo being the best of STEP-xx).

**Figure 4.**
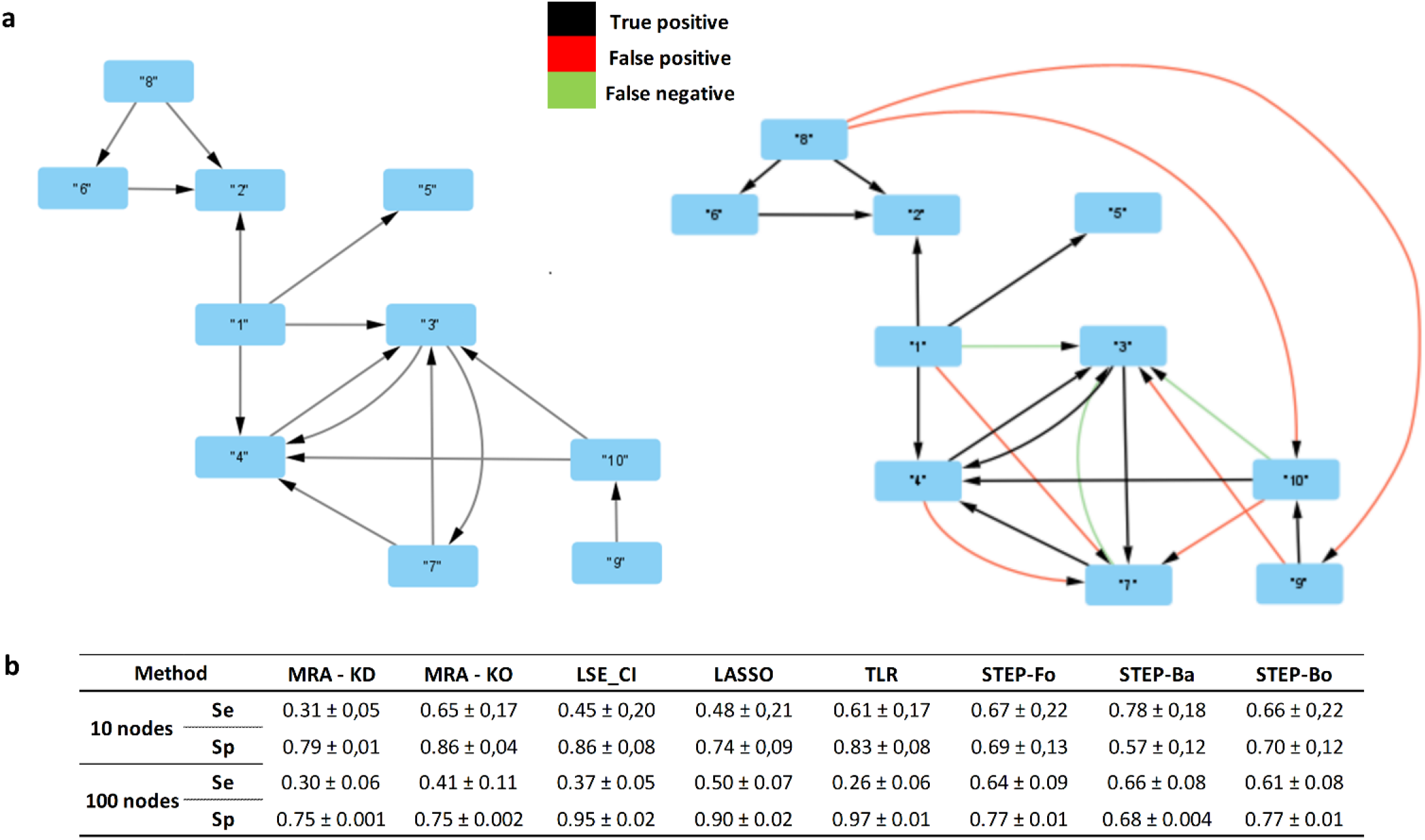
Dream 4 Challenge networks. (**a**) A 10-gene network topology (left) compared to TLR inference (right). (**b**) Average performance of the methods over the five 10- and five 100-gene networks. Se = sensitivity and Sp = specificity.

Naturally, these rankings do not take into account methods developed since DREAM 4. In particular, a variation of MRA called MLMSMRA (Klinger and Blüthgen, 2018), using a likelihood estimation combined with a greedy hill-climbing model selection approach, ranked 3^rd^ with 10-networks and KO perturbations, but 25^th^ with KD. The authors did not report performance for 100-node networks. A Bayesian-modified version of MRA was proposed to better model sparse networks, according to another ranking system defined by the authors themselves (Santra *et al*., 2013; Halasz *et al*., 2016; Klinger and Blüthgen, 2018), *i*.*e*., achieving a performance comparable to MRA combined with STEP-xx.

### The use of *a priori* knowledge of the network topology

In practice, it is common to know about some interactions in a studied network. Regarding known edges, there is nothing particular to do since MRA formalism considers all the interactions as possible. On the other hand, by knowing impossible interactions, we can eliminate equations (see Methods). This increases over determination of the linear system of the regression approach, which is expected to increase the accuracy of the estimates. For methods such as LASSO that require hyper-parameter adjustment through CV, an excess of equations has the potential to improve the optimality of those hyper-parameters.

In the 10- and 100-gene networks of the DREAM 4 Challenge, we assumed different percentages of known null edges. The impact of this knowledge on sensibility and specificity for three of the studied regression methods (LASSO, STEP-Fo, and TLR) is featured in Fig. 5. Indeed, performance improved as a function of the percentage of known null edges. This improvement was stronger for specificity. We observed high variability of the impact on sensibility for 10-gene networks, results were more stable for 100-gene networks.

**Figure 5.**
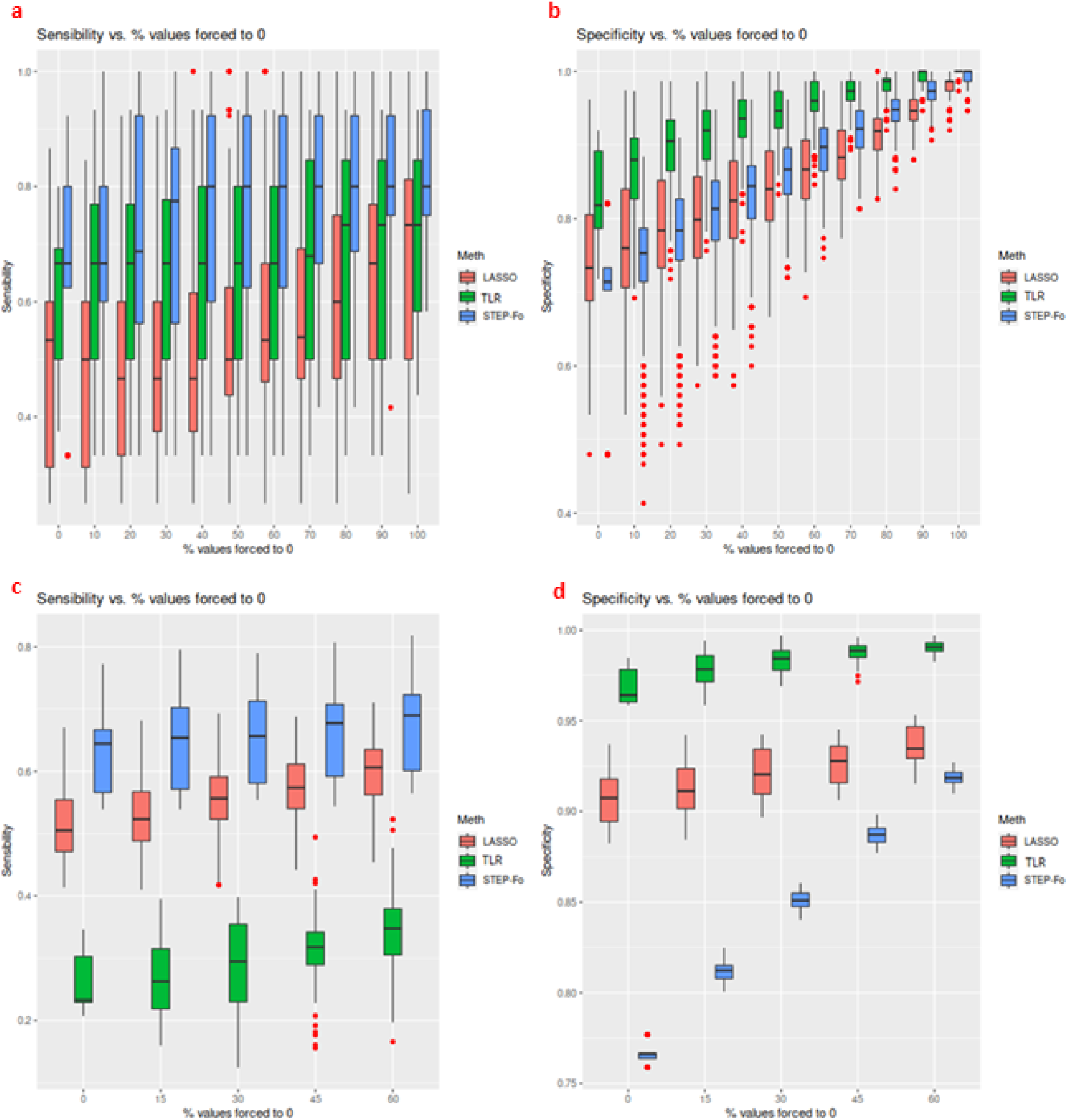
Existing knowledge impact on regression methods tested on DREAM 4 Challenge networks (10 nodes Figs. 5a and 5b, 100 nodes Figs. 5c and 5d). We report the method specificities and sensibilities depending on the number of known null edges.

In comparison with the binary approach proposed here, a Bayesian implementation of MRA has been proposed (Santra *et al*., 2013) that took a known pathway topology as a prior and showed improved accuracy over pure MRA. This Bayesian formulation was less radical than imposing null edges since an absent edge of the pathway could be eventually inferred *a posteriori*, provided that sufficient data evidence existed. On the other hand, our regression-based solution provides accurate CI estimates that hence enable pruning network edges based on model accuracy. One can imagine following with a reduced model, where pruned edges would be removed as above, to obtain better estimates on the other edges. This would be different from a Bayesian formulation obviously, though it would provide comparable practical value.

In the case of multilinear regression, the integration of the knowledge of null edges is done naturally (overfitting the system by deleting columns). In the Bayesian approach (Santra *et al*., 2013), the integration of this knowledge is also very natural (defining the prior probability of the Boolean variables associated with the existence of edges). In MLMSMRA (Klinger and Blüthgen, 2018), the integration of prior knowledge was not explicitly discussed. Nonetheless, because their algorithm fills the connectivity matrix iteratively, it would be easy to force certain coefficients to remain at zero.

## Conclusion

We have introduced a new method to solve MRA equations through multilinear regression. This formulation brings a number of advantages over the classical approach by providing a natural way to model data variability across experimental replicates, or even multiple perturbations at certain or all the modules. Better estimates of MRA connectivity coefficients can be exploited to identify absent edges in a biological network more accurately. Moreover, these advantages were obtained within MRA formalism that provides an elegant, physical interpretation of connectivity coefficients compared to purely regressive approaches.

While motivating the use of regression for MRA, we conducted an analysis of the relationship between data noise, perturbation intensity, and MRA result accuracy that has interest *per se*. In agreement with the local development in Taylor series, MRA accuracy decreases with stronger perturbations as soon as the system is nonlinear. On the other hand, the presence of noise requires perturbations with a minimal intensity to obtain exploitable differences in the variables. Altogether, this means that a compromise must be found between noise levels and perturbation strengths. Depending on the biological system at hand, *i*.*e*., on the nonlinearities, this may lead to perturbations intensity no longer compatible with MRA linear approximation. In such a case, another modeling paradigm must be chosen. In the absence of strong nonlinearities, our work dramatically extended the domain of application of MRA to much larger networks of sizes up to 100. This is a 10-fold increase compared to MRA with standard linear algebra, which had difficulties going beyond 10- node networks in our experiments.

Finally, the proposed approach actually defines a family of MRA-derived methods with the multilinear regression algorithm as a free parameter. While LSE_CI for instance selects edges based on a direct exploitation of residual variance CIs, other algorithms perform model selection such as LASSO or STEP-xx. There are many other regression methods that we have not tested, which might provide specific advantages depending on the dataset, or the specific interest or requirements of the researchers.

