## SUPPLEMENTARY INFORMATION for "Modular response analysis reformulated as a multilinear regression problem"

**1/ A 6-Node Model of MAP kinases**

This model was introduced in Sontag et al. (Kholodenko *et al.*, 2002)

Concentrations and the Michaelis-Menten constants (K_ij_, i= 1, 3, 5, 7, 9, 11; j= 1, 2, 3; K_mp_; K_i_) as well as the catalytic rate constants (k_i_^cat^ , i= 1, 2, 5, 6, 9, 10) and the maximal enzyme rates (V_i_^max^, i= 3, 4, 7, 8, 11, 12) are provided by the authors in their Supplementary Material.

The kinetic equations and moiety conservations derived from the stoichiometry are the following:

d[MKKK-P]/dt = *v*_1_ - *v*_2_ + *v*_3_ – *v*_4_;

d[MKKK-PP]/dt = *v*_2_-*v*_3_;

d[MKK-P]/dt = *v*_5_ – *v*_6_ + *v*_7_ – *v*_8_;

d[MKK-PP]/dt = *v*_6_ – *v*_7_;

d[MAPK-P]/dt = *v*_9_ – *v*_10_ + *v*_11_ – *v*_12_;

d[MAPK-PP]/dt = *v*_10_ – *v*_11_;

[MKKK]_total_ = [MKKK] + [MKKK-P] + [MKKK-PP];

[MKK]_total_ = [MKK] + [MKK-P] + [MKK-PP];

[MAPK]_total_ = [MAPK] + [MAPK-P] + [MAPK-PP].

By inserting the conservation equations in the differential equations, we obtained a system of 6 differential equations that was integrated numerically with the ode R package. The values of the exact connectivity coefficients were computed based on the numerical approximations using the formula

$r_{ij}^{e}=-\left( \frac{X_{j}\frac{\partial f_{i}}{\partial X_{j}}}{X_{i}\frac{\partial f_{i}}{\partial X_{i}}} \right)$.

To determine a time $t_{0}=1000$[s] after which steady state was reached, we used R Deriv library to compute an expression for $\dot{X}(t)$ and found its roots with the library multiroot (one real-valued root was found only). We validated numerically that for $t>t_{0}$ the solution $X(t)$ remained constant.

**2/ Supplementary Tables & Figures**

**k = 0.1%**

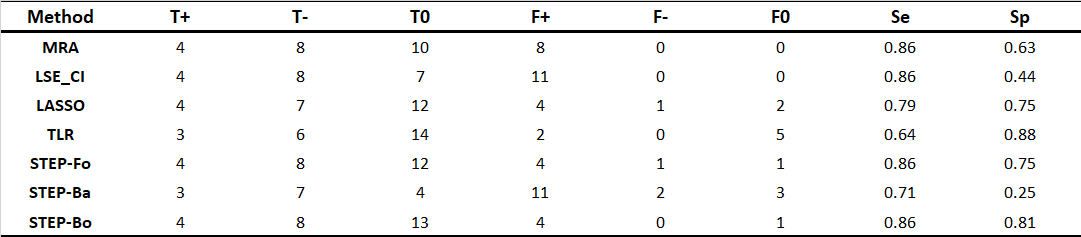

**k = 0.5%**

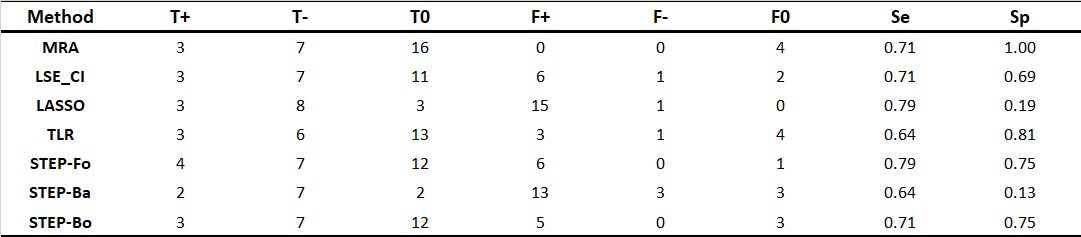

**k = 1%**

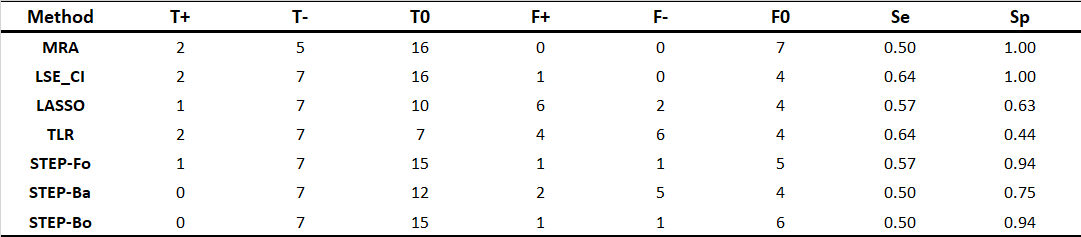

**Supplementary Table 1.** Confusion matrices for the 6 MAP kinase network and three levels of noise. T+ stands for true (correct) prediction of a positive coefficient, T- true prediction of a negative coefficient, T0 true prediction of a zero coefficient, F- is a predicted negative coefficient although its true value is positive or null, etc. Se stands for sensibility and Sp for specificity.

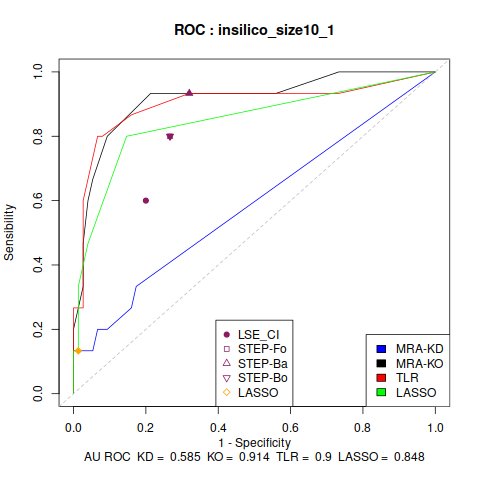

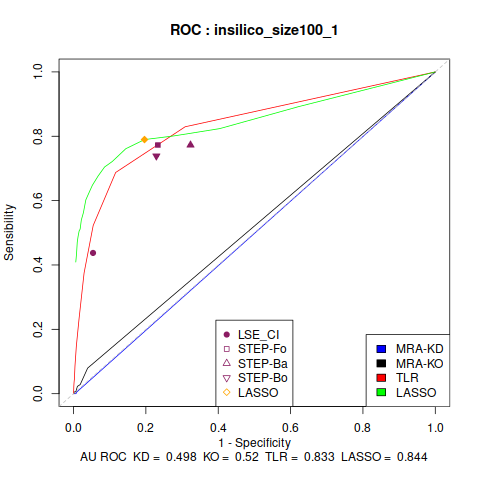

**Supplementary Figure 1.** ROC curves for DREAM 4 Challenge concerning two networks. For other networks ("insilico_size10_x" and "insilico_size100_x"), results are similar to these ones.

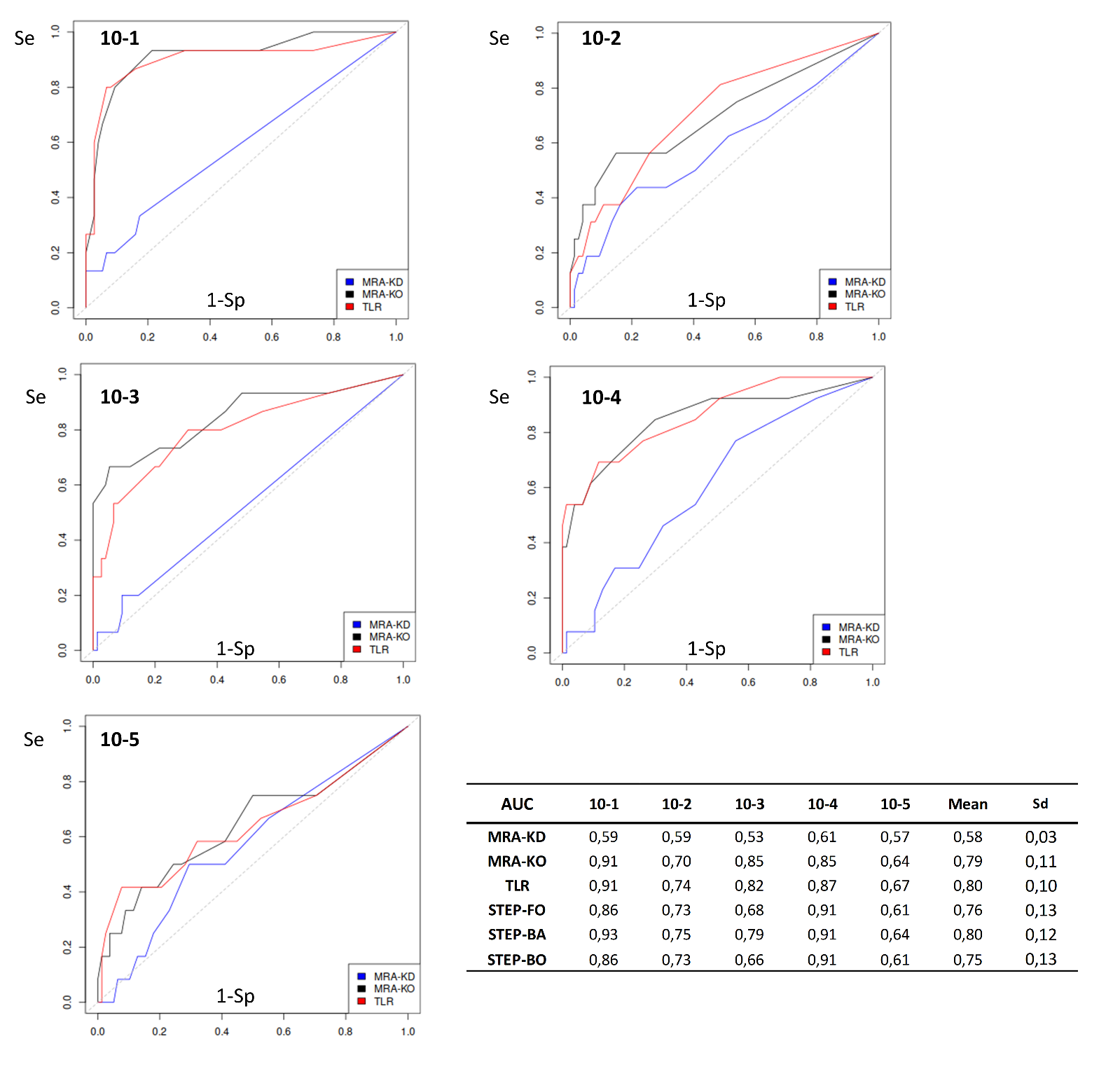

**Supplementary Figure 2.** Representative ROC curves and all the areas under the ROC curve for each DREAM 4 Challenge 10-node network.

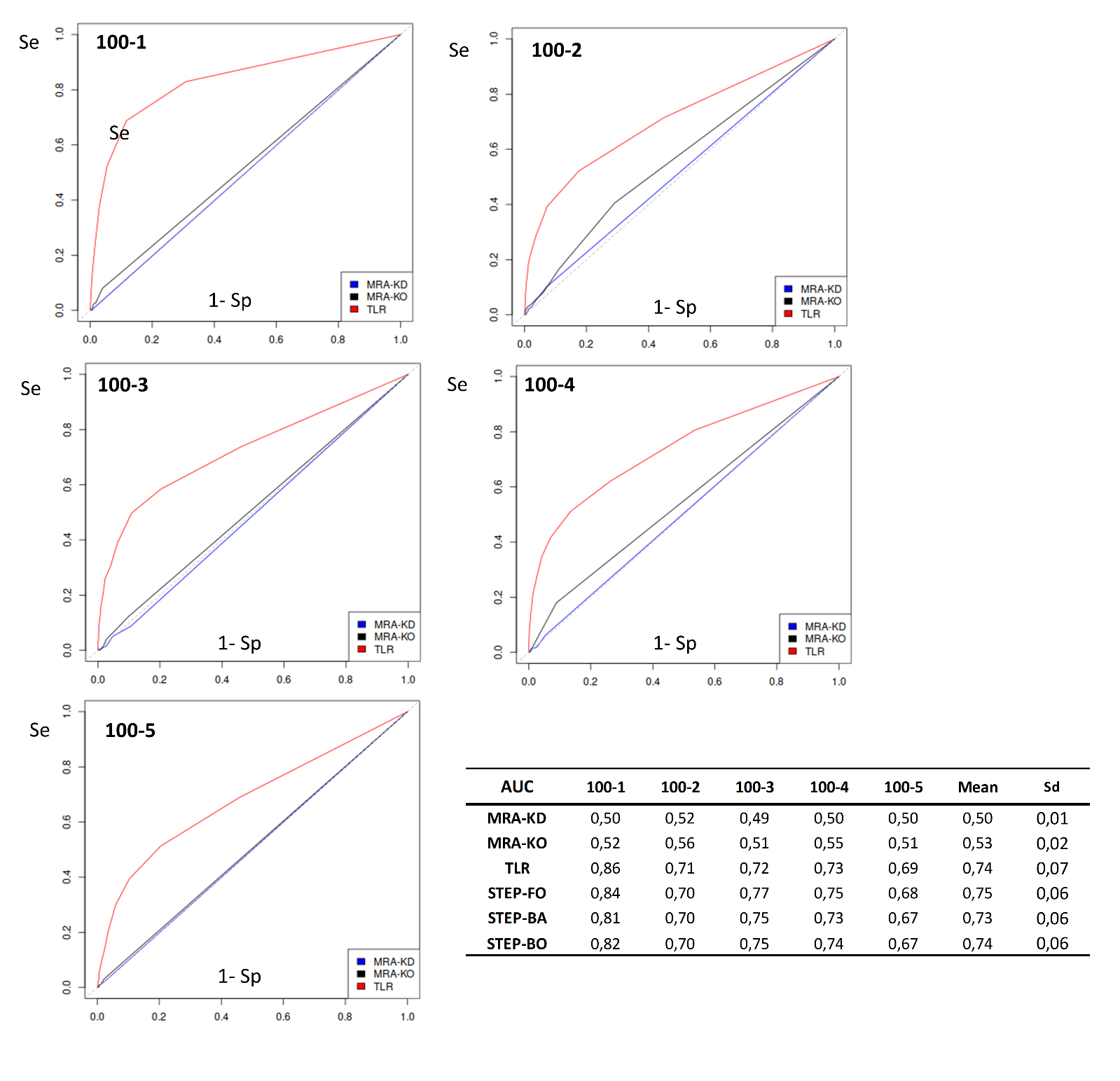

**Supplementary Figure 3.** Representative ROC curves and all the areas under the ROC curve for each DREAM 4 Challenge 100-node network.
